# Effects of cholinergic potentiation on resting-state functional connectivity in healthy adults

**DOI:** 10.64898/2026.09.09.750186

**Authors:** Shea Heaney, Lauryn Foster, Elena Stich, Ioana Grigoras, Charlotte J Stagg, Polytimi Frangou

**Affiliations:** Centre for Integrative Neuroimaging, University of Oxford, Oxford, UK; UKRI MRC Centre of Research Excellence in Restorative Neural Dynamics, UK; Department of Experimental Psychology, University of Oxford, Oxford, UK; Max Planck Institute of Psychiatry, Munich, Germany

## Abstract

Acetylcholine is a key attentional neuromodulator, yet its effects on large-scale organisation of resting-state networks remain poorly understood. Here, we investigated whether acute cholinergic potentiation with the acetylcholinesterase inhibitor donepezil modulates functional connectivity within and between resting-state networks involved in attention. Nineteen healthy young adults completed a double-blind, placebo-controlled crossover study using 7T resting-state functional MRI (rs-fMRI), with imaging performed before and after administration of donepezil or placebo. We tested whether donepezil altered functional connectivity within and between the dorsal attention, ventral attention, default mode, and frontoparietal networks and whether it improved attentional orienting in the same volunteers.

Voxel-wise analysis revealed a significant donepezil-related increase in dorsal attention network connectivity within the left inferior parietal cortex. We did not observe donepezil-related changes within the ventral attention, default mode, or frontoparietal networks. Network-level analyses did not identify significant global changes in network strength. Exploratory analyses indicated reduced coupling between the dorsal attention and default mode networks following donepezil relative to placebo, although this result did not survive correction for multiple comparisons. Donepezil did not significantly improve attentional orienting at the group level. However, exploratory analyses suggested that greater donepezil-related enhancement of functional connectivity within the identified left inferior parietal cluster was associated with greater improvement in attentional orienting.

These findings suggest that acute cholinergic potentiation does not globally enhance resting-state connectivity across attentional networks in healthy young adults. Instead, donepezil may selectively modulate localised dorsal attention network circuitry, particularly within left inferior parietal cortex, a region implicated in sensory integration and externally oriented cognition.

## Introduction

Acetylcholine (ACh) is an important neuromodulator for cognition, with established roles in arousal, sensory processing, cortical plasticity, and attention^1^. Unlike classical fast neurotransmission, neuromodulators such as ACh influence cognition by altering neuronal gain, salience, and attentional state^2^. Cholinergic signalling may therefore provide a mechanism through which attentional state is regulated across distributed brain systems. In particular, cortically-projecting cholinergic neurons of the basal forebrain^3,4^ are well positioned to modulate the large-scale functional networks that support attention. However, how cholinergic potentiation influences the intrinsic organisation of these networks in humans remains poorly understood.

Attentional control depends on coordinated activity within and between several large-scale resting-state networks (RSNs), including the dorsal (DAN) and ventral attention networks (VAN), frontoparietal network (FPN), and default mode network (DMN)^5–8^. The DAN is associated with sustained top-down attention^9–11^, and canonically comprises core nodes in bilateral frontal eye fields (FEF), intraparietal sulcus (IPS), and superior parietal lobule (SPL), while also encompassing a distributed array of other functionally connected brain areas^12^. In contrast, the VAN, comprising right-lateralised temporoparietal junction (TPJ), ventral frontal cortex (VFC), and insular cortex (IC), is more typically associated with stimulus-driven reorienting to unexpected salience^9^. The DAN and VAN are thought to operate in a complementary manner^13,14^, with reduced VAN engagement during focused attention potentially limiting interference from irrelevant stimuli^15,16^. The DMN comprises medial prefrontal cortex (mPFC), posterior cingulate cortex (PCC), medial temporal lobe (MTL), and lateral parietal areas such as the angular gyrus (AG). It is associated with self-referential, mnemonic, and internally-oriented cognition^17,18^ and often shows antagonistic or negative coupling with attentional RSNs^19,20^. This segregation may support externally-oriented cognition by limiting interference from internally-generated processing^21,22^. Finally, the FPN comprises lateral PFC (lPFC) and inferior parietal lobe (IPL) and supports flexible cognitive control^5^, including executive control^23^ and task-switching^24,25^. It may therefore coordinate dynamic interactions between attentional and default-mode systems. Thus, modulating engagement within and interactions between these networks provides a potential large-scale mechanism through which cholinergic signalling could influence attentional state.

Cross-species experimental evidence links cholinergic signalling to attentional processing^26^. Prior work in rodents suggests that cortical ACh release increases during attentionally-demanding tasks^27,28^, while perturbation of cholinergic signalling impairs functional connectivity in attentional circuits^29^ and attentional performance^30,31^. In humans, pharmacological potentiation of cholinergic signalling has similarly been shown to modulate attentional performance^32,33^, while anticholinergic manipulation disrupts selective attention^34^. Pharmacological studies have demonstrated that modulation of cholinergic signalling alters the balance between internally- and externally-oriented cognition. Nicotine has been associated with changes in DMN connectivity^35–37^, while anticholinergic agents alter attentional RSN organisation^37^ and reduce the DMN’s characteristic task-induced suppression^38^. Given the antagonistic relationship between the DAN and DMN during externally oriented cognition^19,20^, these findings raise the possibility that cholinergic potentiation may promote greater functional segregation between these networks. Together, they suggest that cholinergic modulation may influence functional interactions within and between RSNs, but its specific effects on the intrinsic organisation of attentional networks remain poorly understood.

Donepezil (DPZ) is an acetylcholinesterase inhibitor that extends the synaptic availability of ACh^39^ and is currently among the most widely-used symptomatic treatments for cognitive impairment in neurodegenerative disorders^40^. DPZ provides a means of testing the effects of cholinergic potentiation yet, despite its widespread clinical use, relatively little is known about its brain-wide effects on functional RSNs. To our knowledge, only two studies have investigated the effects of DPZ on RSN organisation in healthy young adults. The first examined DPZ in the context of sleep deprivation and reported reduced functional connectivity (FC) within an FPN/executive-control-network-like system^41^. A second study used seed-based rs-fMRI approaches following a chronic two-week dosing regimen and identified reduced FC between a right IPL node and other right FPN regions^42^. However, neither study explicitly interrogated interactions among attentional RSNs nor examined their relationship with behavioural measures of attentional orienting.

Here, we investigated how acute cholinergic potentiation facilitated by DPZ administration alters RSN FC and attentional processing in healthy young adults. Using a double-blind, placebo-controlled crossover design, participants underwent rsfMRI before and after receiving DPZ or placebo (PCB). We focused on the DAN, VAN, DMN, and FPN due to their complementary roles in attentional control and regulating internally- vs. externally-oriented cognition. Given the established involvement of cholinergic signalling in attention^1,26^, we hypothesised that cholinergic potentiation would preferentially modulate FC within the attentional RSNs and their interactions with the DMN. We additionally explored whether DPZ altered coupling between the DAN, VAN and DMN. Based on the complementary roles of the DAN and VAN in externally oriented attention^13–15^, we predicted increased DAN-VAN coupling following DPZ. Conversely, given the antagonistic relationship between DAN and DMN^19,20^, and evidence that cholinergic manipulation alters the balance between externally- and internally-oriented RSN organisation^35–38^, we predicted reduced DAN–DMN coupling following DPZ. Finally, we tested whether DPZ-related changes in attentional RSN FC were associate with changes in attentional orienting. By characterising how acute cholinergic potentiation alters large-scale RSN organisation in healthy cognition, this work may contribute to improved understanding of the role of cholinergic signalling in attentional state regulation and inform development of future therapeutic interventions for cholinergic deficits in neurodegenerative disease.

## Results

### Voxel-wise Effects of DPZ on Resting-State Functional Connectivity

First, we tested whether DPZ altered FC within the DAN, VAN, FPN, and DMN. For each RSN we computed the model fit of each voxel to the RSN’s time series. We subsequently identified voxels showing FC change (post–pre drug administration) that was significantly different following DPZ vs. PCB administration. Statistical significance was assessed using cluster-based inference (cluster-forming threshold *z*=3.1).

Cluster-based inference yielded a significant, 296-voxel cluster in the DAN, localised within left IPL (peak MNI coordinate: x=-54.3, y=-27.5, *z*=41.2, **p_FWE_=0.009**) (Fig. 1). The direction of this effect reflected a greater post-pre FC increase following DPZ compared to PCB (Fig. 1B). No significant voxel-wise effects of DPZ were identified within the VAN, FPN or DMN. We used the Juelich Histological Atlas^43^ to anatomically identify the brain region overlapping with the DAN cluster. The cluster’s peak was localised to tenuicortical supramarginal gyrus (Brodmann area PFt^44^) with 50% probability.

**Figure 1:**
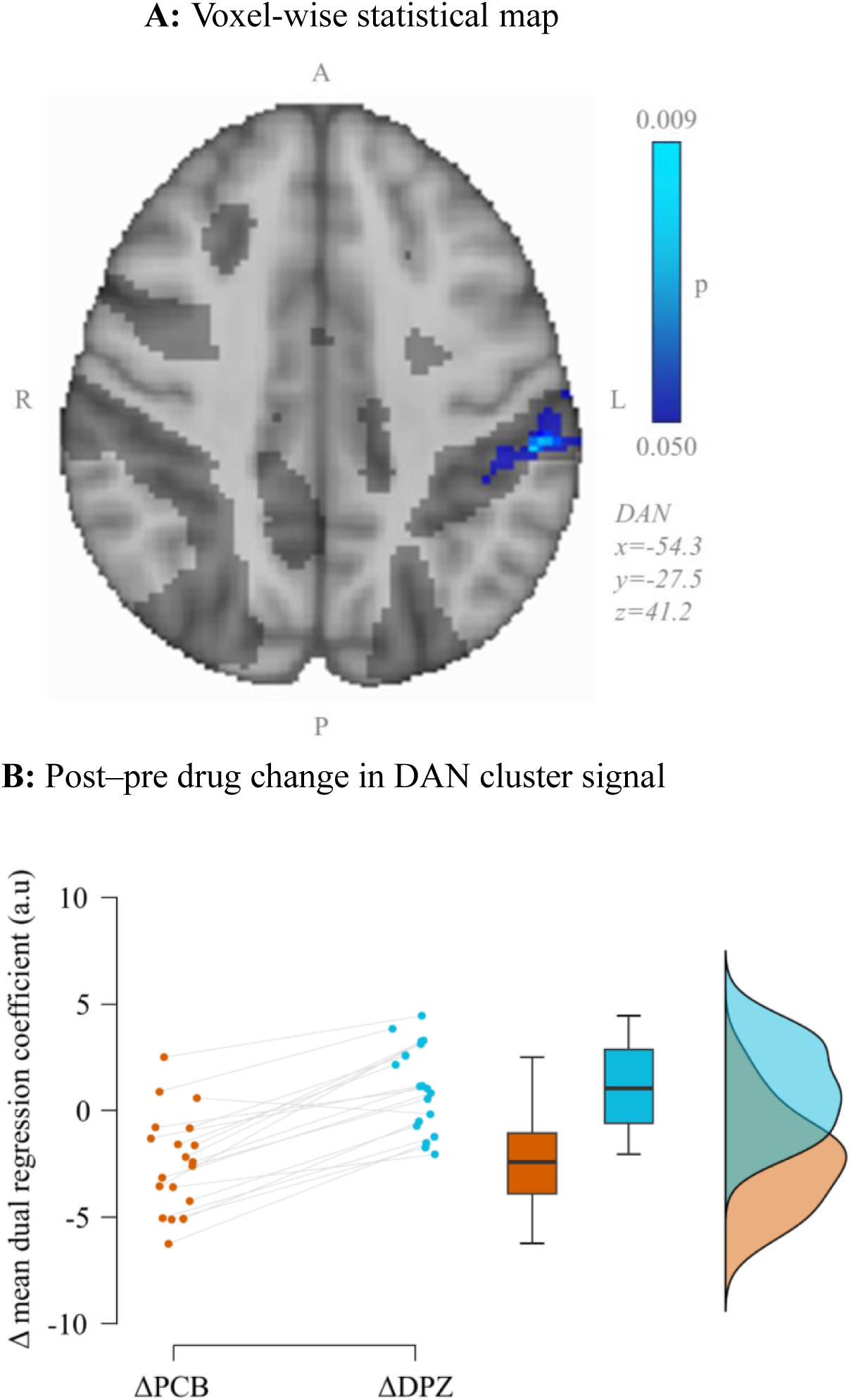
Dual regression analysis yielded a significant cluster in left IPL, whereby changes in functional connectivity following DPZ were significantly different from those following PCB. **A:** A FWE-corrected statistical heatmap of the cluster-wise output, thresholded at p_FWE_<0.05. **B:** A raincloud plot of the difference in mean dual regression coefficients between before (pre-) and after (post-) drug administration (Δ) for placebo (PCB; orange) and donepezil (DPZ; light blue) sessions within the DAN cluster.

### Network-wide effects of DPZ on Resting-State Functional Connectivity

Next, we tested whether DPZ produced more global changes in FC within the four RSNs. For each participant, we calculated network strength as the mean dual regression coefficients extracted from each RSN mask and tested the effects of drug (DPZ vs. PCB), time (pre vs. post-administration) and RSN (DAN, VAN, FPN, DMN) using a linear mixed-effects model (LME) with participant included as a random effect.

There was no significant drug × time interaction (F(1,198.00)=0.259, p=0.611), providing no evidence of an overall effect of DPZ on network strength. The drug × time × RSN interaction was also not significant (F(3,198.00)=2.060, p=0.107), providing no evidence that the effect of DPZ differed across the four RSNs. There was a significant main effect of RSN (F(3,19.59)=16.485, **p<0.001**), but no significant main effect of drug (F(1,18.10)=3.635, p=0.073) or time (F(1,18.23)=0.749, p=0.398).

Planned RSN-specific contrasts similarly identified no significant DPZ-related change in network strength after Holm correction (Fig. 2; DAN: t(70.704)=1.699, p=0.375; DMN: t(70.704)=1.179, p=0.485; FPN: t(70.704)=-1.509, p=0.408; VAN: t(70.704)=-0.334, SE=0.414, p=0.740).

**Figure 2:**
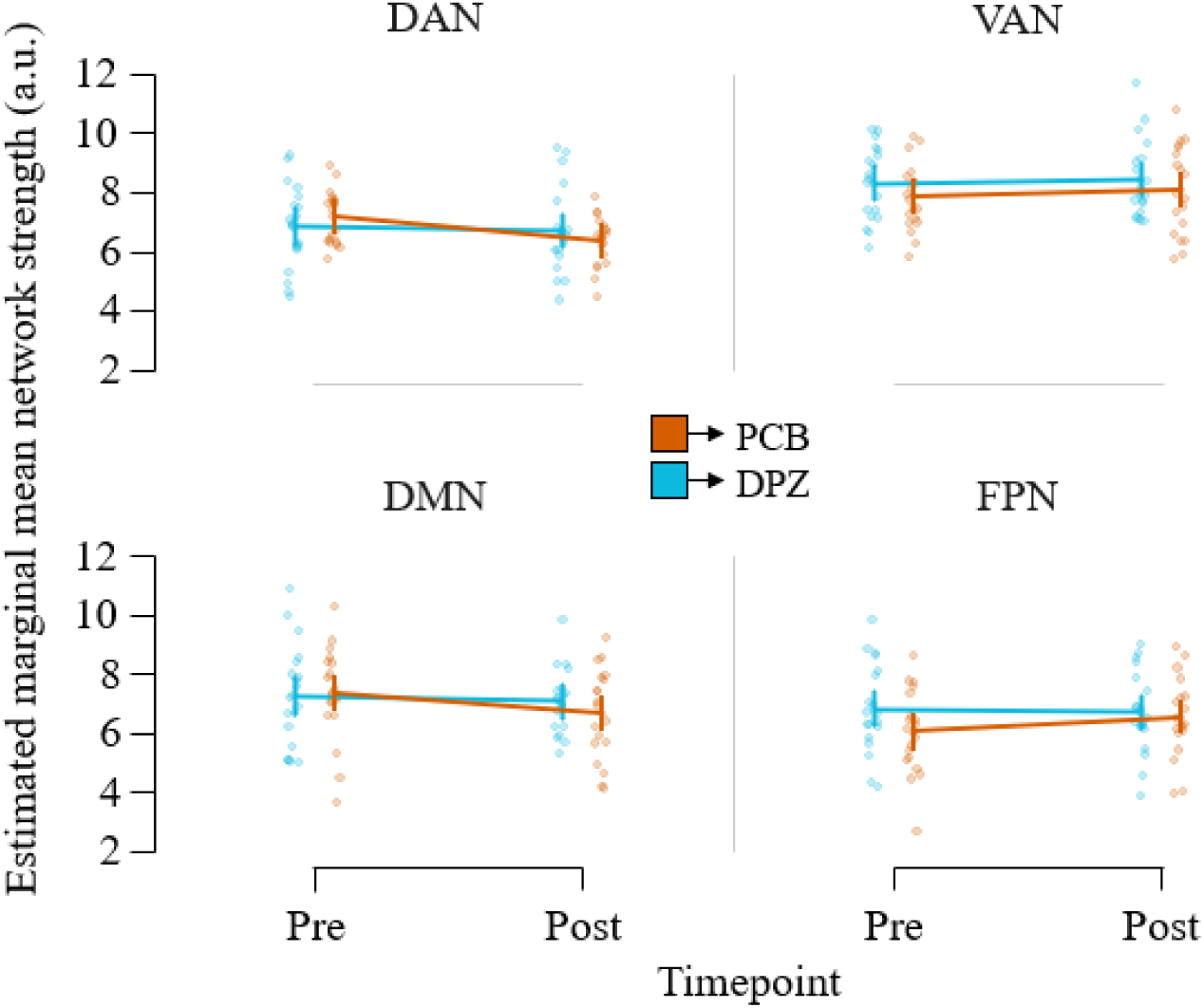
Network strength post–pre change following DPZ vs. PCB. Trajectory plot of the estimated marginal mean and standard error of network strength across pre- and post-drug administration timepoints for the four RSNs under PCB and DPZ conditions. Individual participant network strength points are underlaid.

### Effects of DPZ on RSN Coupling

We next investigated whether DPZ altered FC between the DAN, VAN and DMN. Based on their interactive roles in internally- and externally-oriented attention, we predicted that DAN– VAN coupling would increase following DPZ relative to PCB, whereas DAN–DMN coupling would decrease. For each session and timepoint, inter-network coupling was quantified by correlating RSN time series and applying Fisher’s z transformation. Post–pre coupling changes were then compared between the DPZ and PCB conditions using directional, one-tailed tests.

The change in DAN-VAN coupling did not significantly differ between DPZ and PCB (t(18)=-0.698, p=0.247, d=-0.160; one-tailed; Fig. 3). For DAN–DMN coupling, the post–pre change was lower following DPZ relative to PCB (t(18)=-2.053, **p=0.027**, d=-0.471; one-tailed; Fig. 3), consistent with the predicted direction of reduced DAN-DMN coupling. However, this effect did not survive Holm correction for multiple comparisons _(_p_Holm_=0.054) and should, therefore, be interpreted as exploratory.

**Figure 3:**
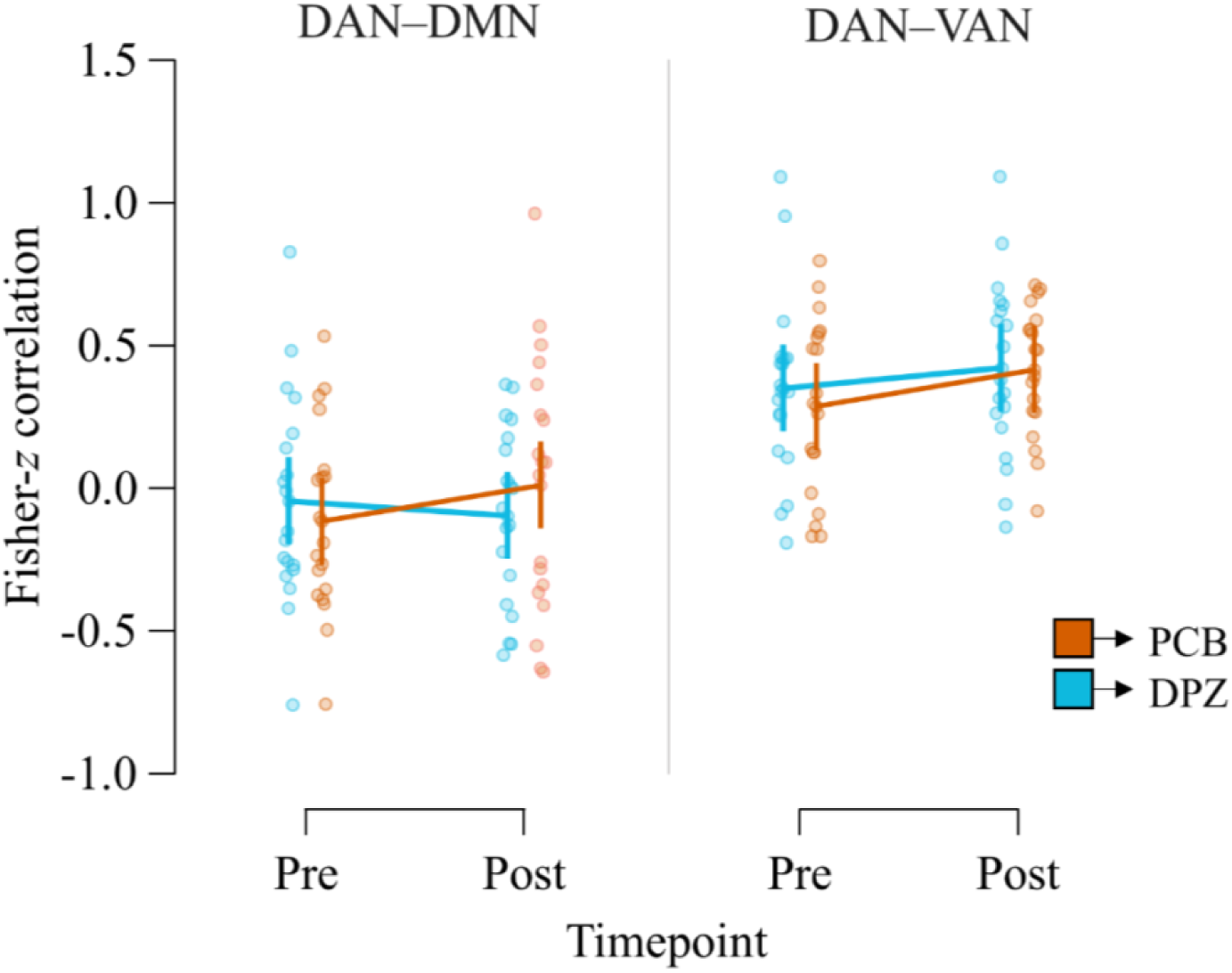
Network coupling. Trajectory plots of the mean and standard error of Fisher-*z* transformed correlation coefficients for each network pair, pre- and post-drug administration for the PCB (orange) and DPZ (blue) session. Coupling was computed from ICA-derived dual regression RSN time series without global signal regression, thus we do not expect canonical DAN and DMN baseline anticorrelation. Individual participant correlations are underlaid.

### Effects of DPZ on orienting attention

We next tested whether DPZ improved attentional orienting, measured using the Attention Network Task (ANT)^45^ following the post-administration MRI scan. Orienting index did not significantly differ between DPZ and PCB sessions (t(16)=-0.241, p=0.813; Fig. 4), providing no evidence for a group-level effect of DPZ on attentional orienting.

**Figure 4:**
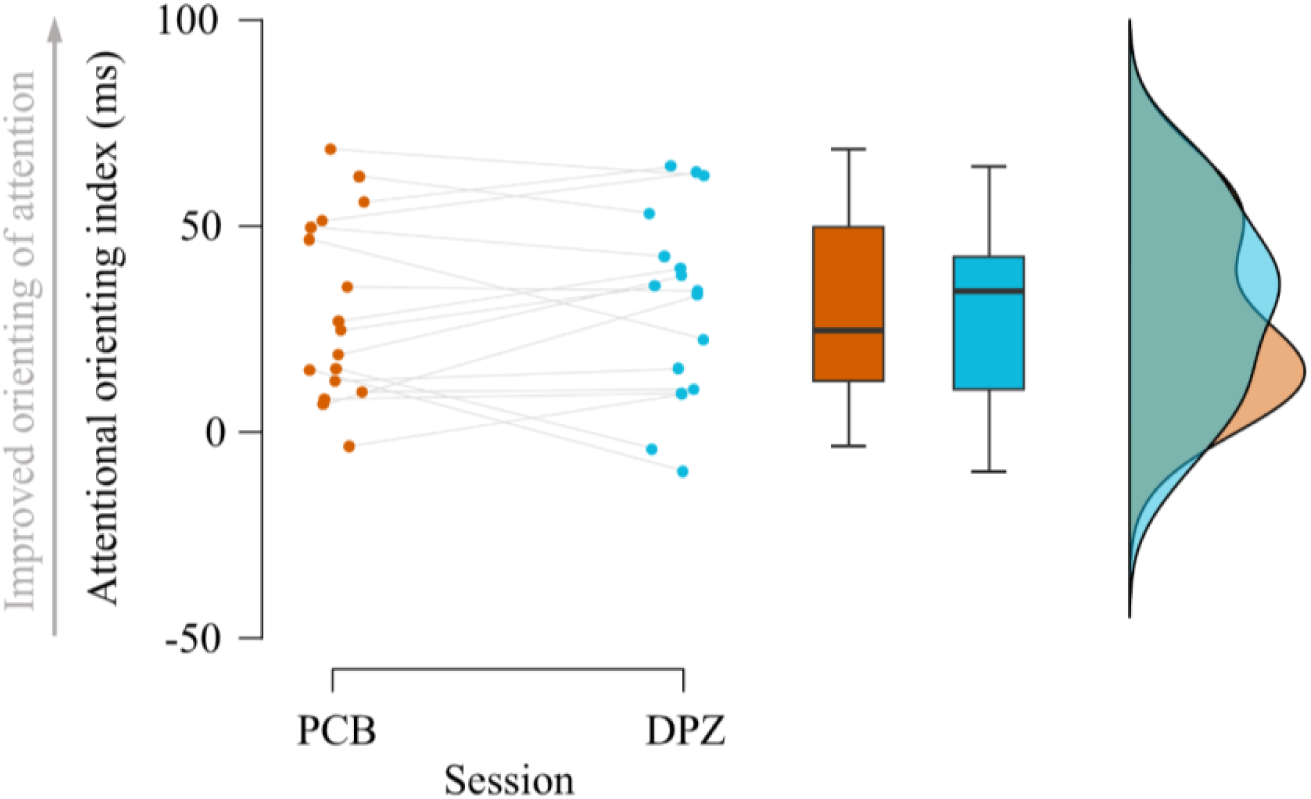
Change in attentional orienting following DPZ vs. PCB. A raincloud plot of attentional orienting index per-participant for PCB (orange) and DPZ (light blue) sessions.

Next, we investigated whether individual differences in the DPZ-related FC change within the identified left IPL/DAN cluster were associated with individual differences in attentional orienting. For each participant, we related the DPZ vs. PCB post-drug difference in FC within the cluster to the DPZ vs. PCB difference in attentional orienting. Greater FC following DPZ compared to PCB was associated with improved attentional orienting in the DPZ session (Spearman’s ϱ=0.431, **p=0.043**; one-tailed; Fig. 5).

**Figure 5:**
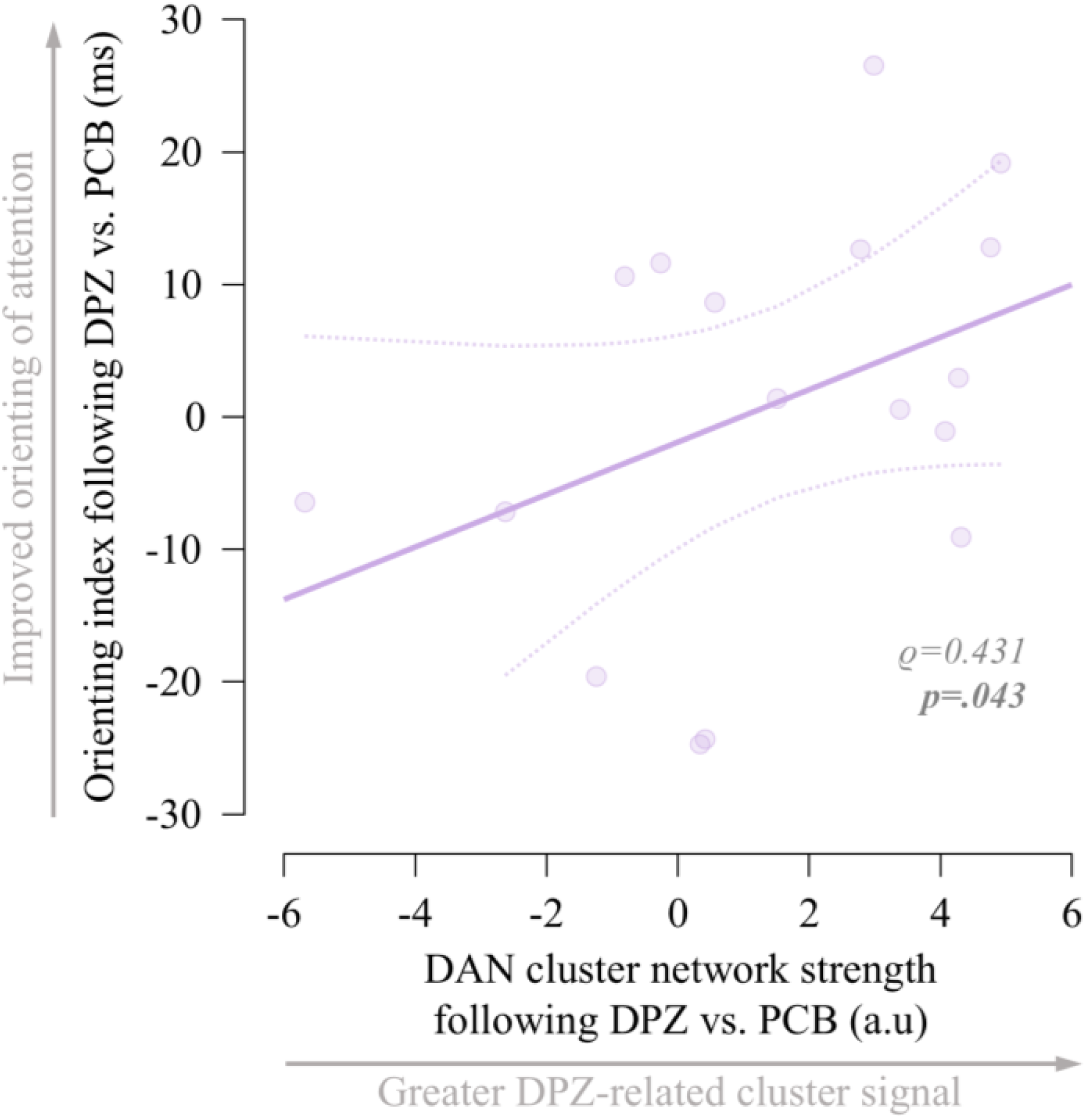
Relationship between DAN cluster network strength and attentional orienting. Scatter plot of the relationship between DPZ-related change in DAN cluster signal and DPZ-related change in attentional orienting index. Index change is computed as the difference between DPZ–PCB sessions. DAN cluster network strength change is computed as the difference between post-drug DPZ–PCB measurements.

## Discussion

In the present study, we investigated the effects of acute cholinergic potentiation on FC within RSNs associated with cognition and attention using a double-blind, placebo-controlled, cross-over rs-fMRI design. DPZ did not produce widespread changes in RSN organisation, but selectively increased FC within the DAN. We found no evidence for network-wide changes in FC, while exploratory analyses suggested that DPZ may also reduce DAN–DMN coupling and that individual differences in FC within the DAN may relate to attentional orienting. Together, these findings suggest that acute cholinergic potentiation may produce selective rather than widespread changes in attentional RSN organisation.

### Donepezil selectively increased localised connectivity within the DAN

Our primary hypothesis was that cholinergic potentiation would alter within-network connectivity of the RSNs involved in attentional processing. Consistent with this hypothesis, we identified a cluster of voxels within the DAN where FC increased following DPZ compared to placebo. This cluster was located within the left IPL and overlapped with tenuicortical supramarginal gyrus (Brodmann area PFt).

The location of the cluster is notable, as it lies more ventrally within the IPL than canonical DAN regions such as IPS and FEF^14^. However, the IPL is functionally heterogeneous, and area PFt has been implicated in higher-order sensorimotor and attentional integration^46–48^, with functional coupling to canonical DAN structures^49^. The present findings therefore suggest that cholinergic potentiation may selectively modulate FC within an attentional integration region of the DAN rather than uniformly increasing connectivity across the entire network. Consistent with this interpretation, we found no evidence of a network-wide change in DAN FC. We found no significant voxel-wise effects of DPZ within the VAN, DMN or FPN and no evidence for network-wide FC changes across the four RSNs.

Together, these findings suggest that acute cholinergic potentiation does not produce widespread changes in FC within attentional networks, but rather selective modulation of DAN-associated IPL, consistent with broader theories proposing that ACh supports externally-oriented attentional processing rather than producing indiscriminate cortical arousal^1^.

### Effects of donepezil on attentional network coupling

We additionally investigated whether DPZ altered coupling between attentional networks and the DMN. DAN–VAN coupling did not significantly differ between DPZ and PCB, whereas the post–pre FC change in DAN–DMN coupling was reduced following DPZ compared to PCB, consistent with our prediction. However, this finding did not survive multiple comparison correction, and should, therefore, be interpreted with caution.

The DAN and DMN are considered antagonistic systems with a relationship often characterised by negative coupling or anticorrelation between the two at rest^19^, although the magnitude of this relationship depends strongly on preprocessing and analytical approach^6^. Literature has shown that increased attentional demand drives further segregation of task-positive systems from the DMN^50^; an effect that is blunted in clinical cases of cholinergic deficit^51,52^. Therefore, our findings are consistent with the idea that cholinergic potentiation may be enhancing functional segregation between externally-oriented attentional systems and internally-oriented default-mode processing. However, given the exploratory nature of the analysis and lack of significance following multiple-comparison correction, this interpretation requires confirmation in a larger independent sample.

### Effects of donepezil on attentional orienting and the relationship with DAN FC

Contrary to our hypothesis, DPZ did not significantly improve attentional orienting at the group level. This may reflect the relatively subtle effects of acute cholinergic potentiation on attentional performance in healthy young adults.

We next explored whether individual differences in FC following DPZ compared to PCB within the left IPL/area PFt cluster were associated with individual differences in attentional orienting following DPZ vs. PCB. Greater DPZ-related FC increases within the left IPL/area PFt cluster were associated with greater improvement in attentional orienting. Although exploratory, this relationship suggests that the effects of cholinergic potentiation on RSN organisation may be behaviourally relevant even in the absence of group-level behavioural effects.

The absence of a group-level behavioural effect may also reflect the sensitivity of the behavioural measure. Prior work has suggested that the classical ANT may be less sensitive to cholinergic modulation than tasks including invalid spatial cues^37,53^. Future studies employing more targeted measures for attentional systems mediated by cholinergic signalling may be better positioned to test the effects of pharmacological interventions.

### Limitations and methodological considerations

Several features of the study should be considered when interpreting these findings. First, we investigated the effects of a single acute dose of DPZ in healthy young adults. Evidence from both cognitive and neuroimaging studies suggests that cholinergic enhancement in healthy populations is subtle and state-dependent^1,54–56^. Unlike clinical populations with existing cholinergic deficit, healthy young adults may already lie near the optimal cholinergic tone, thereby reducing the achievable enhancement from increased cholinergic synaptic availability.

Second, participants viewed nature documentaries during the rsfMRI measurements rather than resting with their eyes open at fixation. Although FC was calculated using a resting-state FC pipeline, the experimental state is best described as naturalistic viewing rather than unconstrained rest. This approach was chosen to support participant tolerability and stable engagement during the lengthy imaging session, but naturalistic viewing itself may engage distributed attentional and sensory systems^57^. Consequently, the present findings may reflect modulation of sustained externally-oriented engagement rather than resting-state organisation; this context may have contributed to the localisation of the FC change within the DAN.

Finally, the modest sample size may have limited sensitivity to smaller network-level effects. The within-subject crossover design reduced between-subject variability^58^, thus increasing sensitivity to pharmacological effects, but does not overcome the limitations associated with a small sample. Accordingly, the absence of significant effects within the other RSNs should not be interpreted as evidence that DPZ has no influence on their functional connectivity. Replication in a larger independent sample will be important, particularly for the exploratory DAN–DMN coupling and brain–behaviour findings.

## Conclusion

Here, we show that acute cholinergic potentiation with DPZ produces a localised increase in DAN-associated FC in healthy young adults, without evidence for widespread changes across the attentional networks. Exploratory evidence suggests that individual variability in this brain response may relate to variability in attentional orienting. These findings provide a basis for further investigation of how cholinergic signalling shapes large-scale attentional network organisation in health and disease.

## Materials and Methods

### Participants

We recruited 20 (12 female, 8 male) healthy young adults (mean age ± SD: 26.33 ± 4.27 years) with normal or corrected-to-normal vision. Exclusion criteria included ongoing use of long-term medication, pregnancy, breastfeeding, any history of or current psychiatric or neurological conditions, diabetes, alcohol or sedative drug abuse, asthma, chronic obstructive pulmonary disease, sick sinus syndrome, supraventricular conduction abnormalities, susceptibility to peptic ulcers, cardiac disease, gastro-intestinal complications, prolactinoma, long QT syndrome, and failure to meet requirements of a 7T MRI safety screening form.

Participant body mass index (BMI) was between 20.31kgm^-2^ – 33.67kgm^-2^ (mean BMI ± SD: 25.39 ± 3.17 kgm^-2^), in line with previous studies probing cognition using DPZ^59,60^. Participant handedness was assessed using the Edinburgh Handedness Inventory^61^ and tasks performed using their dominant hand (right). One participant did not complete all scans due to feeling unwell following drug administration and thus was excluded from imaging analysis (analysed sample n=19; 11 female, 8 male).

Written, informed consent was obtained from participants in accordance with study approval from the Central University Research Ethics Committee (CUREC) (MS IDREC reference: R75163/RE004).

### Pharmacological Intervention

Cholinergic potentiation was facilitated with a 10mg dose of DPZ. DPZ is a centrally-acting, non-competitive, reversible inhibitor of cholinesterases, primarily acetylcholinesterase. Peak plasma concentration is reached at approximately 3-5hrs post administration^62^, with a half-life of approximately 70hrs. By inhibiting acetylcholinesterase, DPZ increases ACh synaptic availability, thereby increasing central cholinergic signalling^39^.

To mitigate potential common side-effects of DPZ due to its peripheral effects (i.e., nausea, vomiting), we administered 10mg of the antiemetic domperidone, simultaneously with DPZ/placebo. Domperidone is a peripherally-acting selective antagonist of dopamine D_2_/D_3_ receptors with minimal crossing of the blood-brain barrier^63^.

### Study Design

Participants were invited to two sessions in which they received either DPZ or PCB in a double-blind, cross-over design. The order of DPZ and PCB sessions was counterbalanced across participants. Sessions were ≥7 days apart to ensure between-session drug clearance. Height, weight, and basal metabolic rate were collected during the first session.

In each session, participants underwent two MRI brain scans: one before and one after drug administration. Each scan lasted approximately 60min, during which we collected whole-brain resting-state data while participants watched a nature documentary without audio. Immediately following the first scan (t=60min), participants were provided with two capsules — containing either the active drug (2×5mg DPZ) or PCB (2 sucrose pills) — and a domperidone pill. The second scan took place 180min after (t=240min) to allow DPZ to reach peak plasma concentration. Immediately following the second scan (t=300min), participants completed a blinding questionnaire and performed the Attention Network Task (ANT)^45^, an established task for investigating attentional processes. Data on participants’ subjective experience were collected using Positive and Negative Affective Schedule^64^ and Bond and Lader Visual Analogue Scale^65^ questionnaires provided at drug administration (t=60min), when drug reached peak plasma concentration (t=240min) and at the end of each session (t=300min). A session outline is illustrated in Fig. 6.

**Figure 6:**
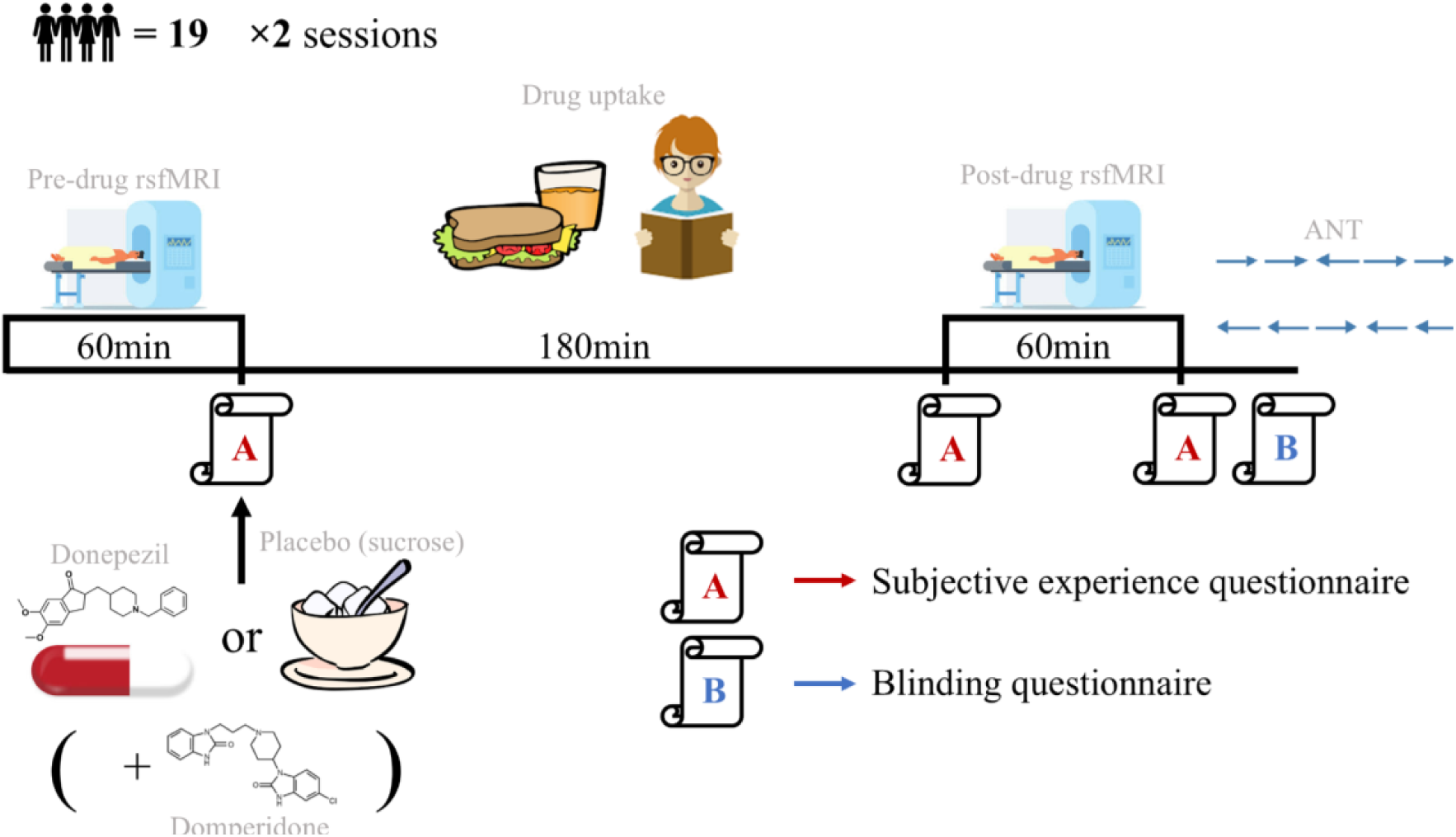
Session outline. A typical session outline. Sessions were identical aside from administered drug condition.

### Set-up

Participants were scanned in a MAGNETOM Investigational Device 7T Plus (Siemens Healthineers, Erlangen, Germany) system using a 32-channel head coil (Siemens Healthineers). Physiological signals – i.e., breathing/heart rate – were collected using BIOPAC respiratory bellows (BIOPAC Systems, Goleta, California, USA) and a BIOPAC pulse meter (BIOPAC Systems) throughout scans. Physiological recordings were collected for monitoring purposes but not explicitly included as regressors. Structured physiological noise was instead addressed through ICA denoising^66,67^.

The ANT was run on a Dell Windows 10 laptop using a publicly-available Python script (pypi.org/project/psychometric-tests). Participants were seated 60cm from the screen and responded using the laptop’s keyboard.

### MRI data acquisition

T1-weighted structural images were acquired using a magnetization prepared rapid acquisition gradient echo (MPRAGE) sequence (TR 2600.0ms, TE 3.18ms, TI 1100ms, TA 3.7min, 1.0mm isotropic, FoV 224×224×176mm^3^, sagittal, PAT factor 3, no fat suppression, flip angle 5.0°). B0 field map magnitude and phase were acquired using a Gradient-Echo (GRE) sequence (TR 620.0mm, TE-1 4.08ms, TE-2 5.1ms, TA 2.0min, 2.0mm isotropic, FoV 192×192×154mm^3^, flip angle 39°). Whole-brain resting-state functional images were collected using a multiband accelerated echo-planar imaging 2D (MBEP2D) sequence (TR 1165ms, TE 20.00ms, TA 5.8 min, 2.0mm isotropic, FoV 220×220×144mm^3^, 287 volumes, multiband acceleration factor 3, flip angle 60°).

### MRI data preprocessing

All imaging data were processed using FMRIB Software Library (FSL^68,69^). Our resting-state pre-processing and analysis pipeline is outlined in Fig. 7.

**Figure 7:**
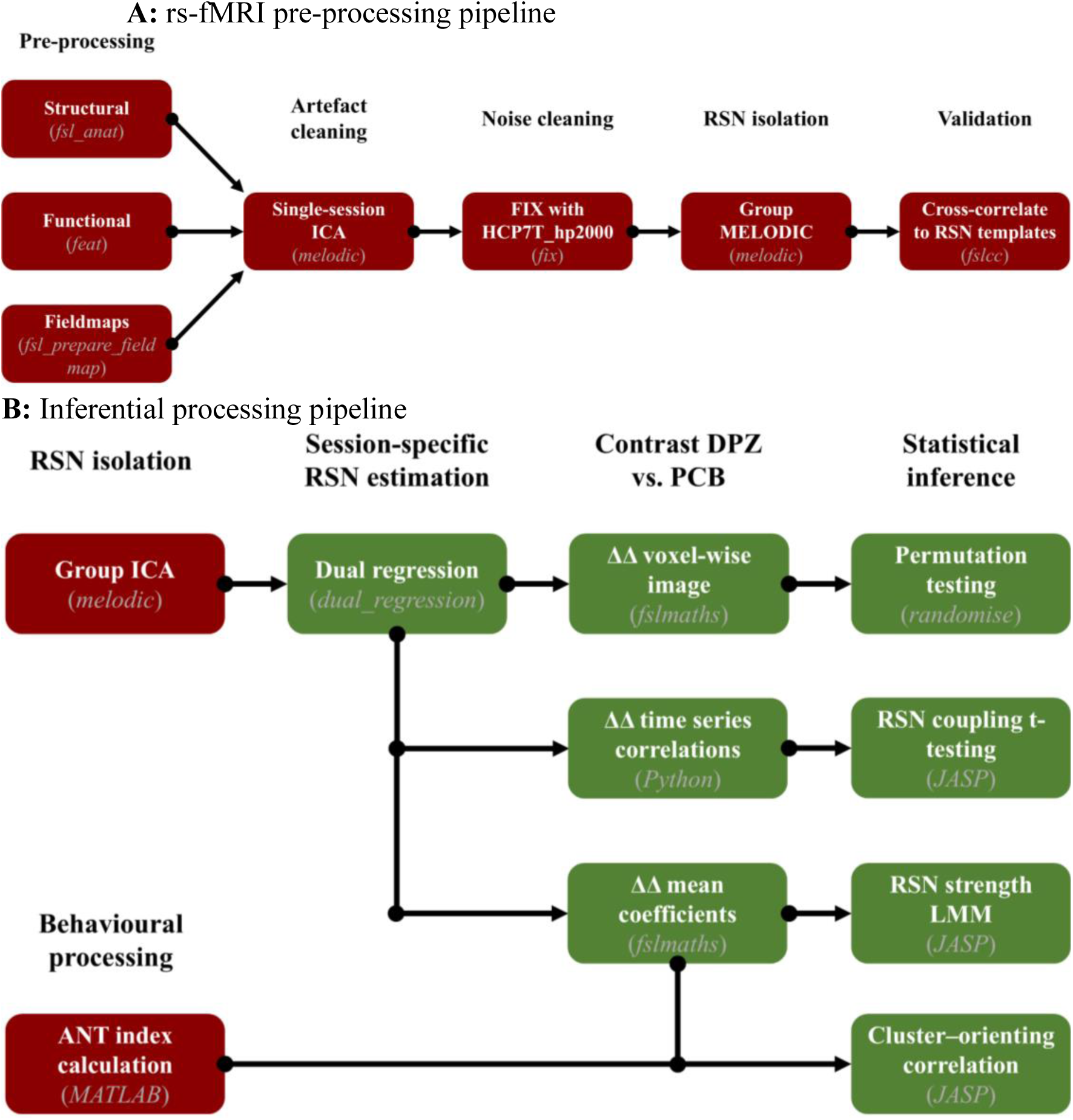
**A:** rs-fMRI preprocessing pipeline. **B:** Inferential analysis pipeline showing derivation of voxelwise ΔΔ images, RSN strength measures, inter-network coupling metrics, and behavioural correlations.

Structural images were pre-processed using the *fsl_anat* pipeline, including bias-correction (*fast*^70^) and brain-extraction (*bet*^71^), with non-linear transformations from participant- to Montreal Neurological Institute (MNI) standard-space (*fslreorient2std*, *flirt*^72–74^, *fnirt*^75^).

Functional (T2*-weighted) images were preprocessed using an adapted *fsl_anat* pipeline, which included motion-correction (*mcflirt*^72^) and bias-correction. To reduce MRI distortions, B0 unwarping was applied to the functional images using the preprocessed field maps. Distortion-corrected images were high-pass filtered at a 100s period cutoff (i.e., <0.01Hz oscillatory activity removed). No smoothing was applied during *feat* preprocessing. Using FSL *feat*^76^, functional images underwent boundary-based registration (BBR) to participant structural-space using transformations generated by *fsl_anat*, then structural-to-standard transformations were applied for standard-space registration.

### Resting-state data analysis

#### ICA Denoising

To remove substantial artefacts, e.g. due to participant motion or MRI-related noise, independent component analysis (ICA) was performed for each timepoint separately using *melodic*^76^. Components were classified using FSL *fix*^66,67^ trained on a standard model set of minimally-processed 7T images from healthy controls (HCP7T_hp2000). We used the default recommended *fix* classification threshold (20%), which defines the balance between true positive and true negative rate, and visually inspected borderline cases (15-25%) to ensure correct classification; no manual relabelling was deemed necessary. Spatial smoothing was applied after *fix* denoising. Clean images were smoothed with a 5mm full-width half maximum (FWHM) Gaussian kernel using *fslmaths*. Smoothed images were registered to MNI standard-space with FSL *applywarp* using prior *fsl_anat* structural-to-standard transformations.

#### Group ICA and Network Identification

Next, we isolated and identified the RSNs of interest: the DMN, FPN, DAN, and VAN. We iteratively explored different fixed dimensionalities using the following two criteria to decide on the optimal number of components. First, prior work has shown that RSNs are strongly reproducible across studies and participant populations^77,78^, thus for each iteration, we visually compared the derived components to canonical RSNs (Fig. 8). Second, we cross-correlated our components with canonical references^79^ using *fslcc* to obtain a quantitative metric for RSN representation (Tbl. 1). For the RSNs of interest, we found the number of components that produced highest correlation coefficients between the canonical RSNs and the derived components, after removing those with *r*<0.15. Using this iterative process, we decided that the optimal dimensionality for the group ICA was 11. In this dimension, the DAN had spatial correspondence of *r*=0.49 with IC2, while the VAN with IC3 (*r*=0.53), the DMN with IC4 (*r*=0.56) and the FPN with IC7 (*r*=0.53). In all cases, ICs were visually inspected and core network nodes were identified.

**Table 1:**
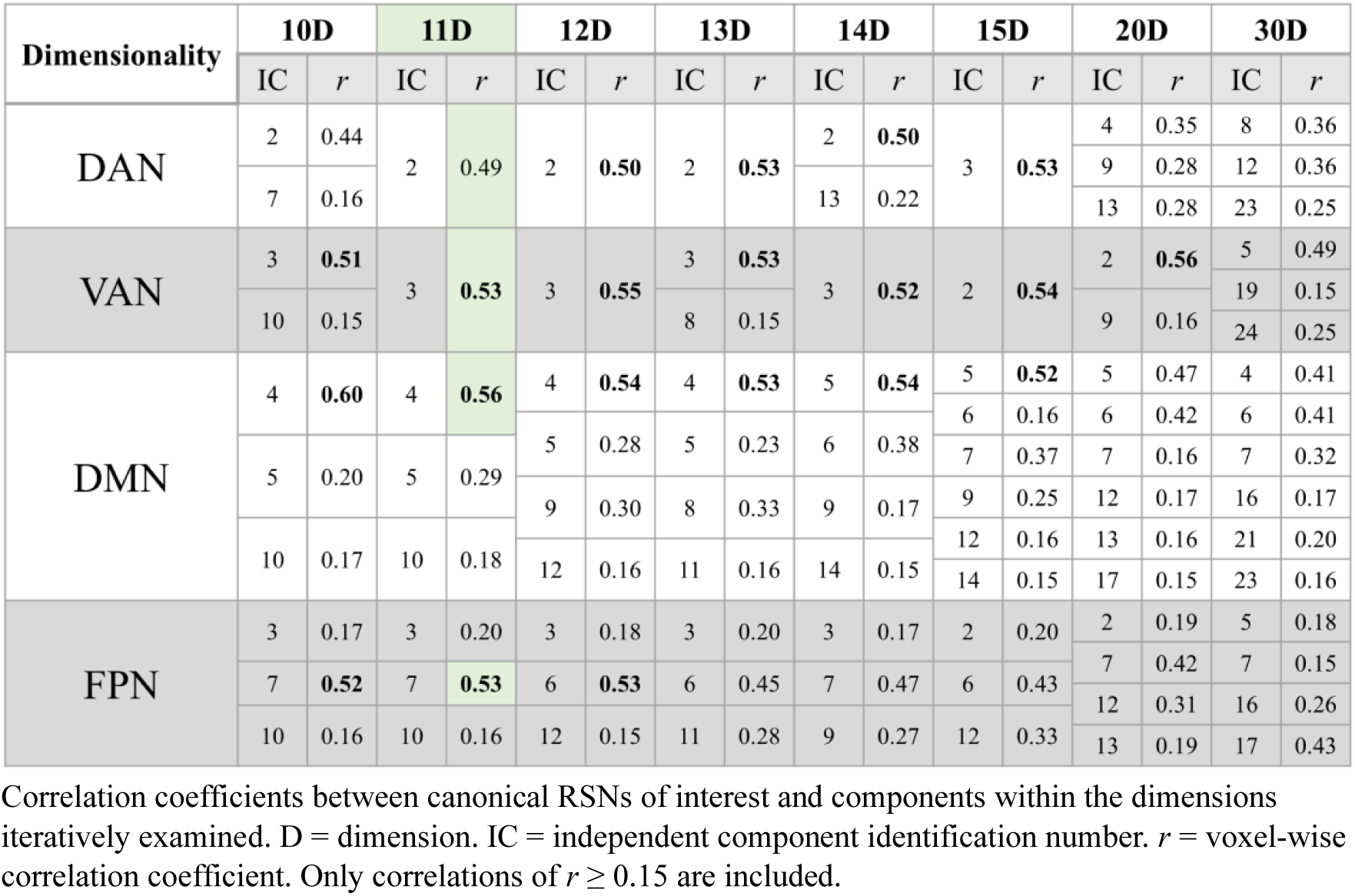
Correlation of canonical RSN templates with examined dimensions.

#### Dual regression

Having identified the RSNs of interest in our participant population, we next tested whether connectivity within these RSNs was increased or decreased by enhanced cholinergic signalling under DPZ. To this end, we obtained participant-, timepoint- and session-specific representations of each RSN using dual regression. Using the FSL tool *dual_regression*^80^, a two-stage analysis was executed to generate 4D spatial RSN maps. First, measurement-specific time series were extracted from the group-level ICA spatial maps. Second, the time series were used to identify matching measurement-specific spatial maps. Ultimately, the maps represent how well each network is expressed in a given scan. The resulting time series and maps for each participant, timepoint (pre-vs. post-administration), and session (DPZ vs. PCB) were used for statistical analyses.

#### Voxel-wise Statistical Analysis

First, we ran voxel-wise statistics. To compute drug-induced connectivity modulations for each voxel within each of the RSNs of interest, we used *fslmaths* to compute an explicit difference-of-differences (ΔΔ) image for each participant and network from the dual regression output, as outlined below. Each participants’ pre-drug maps were subtracted from their corresponding post-drug maps, then, participants’ change following PCB was subtracted from change following DPZ:

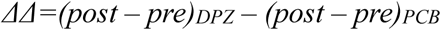

To test for drug-induced modulation, we performed non-parametric permutation testing on the ΔΔ dataset. Here, we used FSL *randomise*^81^ with 5000 sign-flipping permutations to test if ΔΔ ≠ 0. The result was corrected for multiple comparisons using null distribution of the max cluster mass (cluster-wise) correction with a cluster-forming threshold of *z* = 3.1. The analysis output included: raw t-statistic maps, used to estimate effect directionality; and cluster-wise-corrected p-value maps, used to identify significant major clusters. P-value maps were thresholded at p_FWE_<0.05 to identify significant clusters within the RSNs of interest. No covariates were included for this analysis due to homogenous sample characteristics (healthy, young adults) and prior denoising of motion artefacts.

#### Network-level and cluster-based analysis

To quantitatively explore changes in network expression, we used mean dual regression coefficients masked for each network. Each of the four network volumes were extracted from the concatenated *melodic* file, thresholded (*z*>2), and the output binarized to obtain network masks. Using the masks and *fslmeants*, we calculated the mean dual regression coefficient within each RSN mask. These produced scores reflecting the mean degree to which each voxel’s time series was associated with the group-level RSN timecourse for each participant, condition, timepoint, and network, used as a proxy to network strength.

Voxel-wise analysis identified a significant cluster in left inferior parietal lobule (IPL). A brain mask was created by thresholding (p_FWE_<0.05) and then binarising the cluster-wise-corrected map. The mean coefficients were then extracted from the mask.

To interrogate inter-network connectivity, we began by correlating RSN-specific, ICA-derived dual regression stage 1 time series. To this end, the *numpy.corrcoef* function was used for Pearson’s pairwise correlations between DAN–VAN and DAN–DMN time series. This produced an *r* coefficient that was converted to *z*-score using Fisher’s *z* transformation (*z=arctanh(r)*). *z*-scores were subsequently processed as (post – pre)_DPZ_ – (post – pre)_PCB_ to obtain a ΔΔ per RSN pair and per participant.

### Attentional Network Task

In the ANT, participants are asked to indicate the direction of an arrow appearing with flankers at the top- or bottom-half of a computer screen, while maintaining fixation on a central cross. Each trial is preceded by four possible cues (none, central — an asterisk appearing at the centre, double — two asterisks appearing at the top and bottom halves, spatial — an asterisk appearing at the screen half where the arrows will appear) and includes one of three stimulus complexity levels (neutral — the flankers are dashes, congruent — the flankers are arrows pointing in the same direction as the central arrow, incongruent — the flankers are arrows pointing in the opposite direction from the central arrow), in a counterbalanced, randomised order. Participants completed 288 trials in each session, following the post-drug MRI scan. Data were pre-processed (MATLAB R2024a) to remove reaction time (RT) outliers (Grubbs’ test^82^), remove trials without a response, and compute attentional processing indices. Behavioural data from two participants were excluded due to incomplete ANT recordings (analysed sample n=17).

In line with the literature^45^, to capture attentional orienting, we computed the difference in mean participant RT (MRT) following central vs. spatial cue conditions:

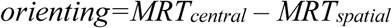

Spatial cues validly predict target location, whereas central cues provide temporal but not spatial information, capturing covert attentional orienting. A greater decrease in RT following spatial cues relative to central cues is indicative of improved attentional orienting.

### Statistical modelling

We explored changes in network strength driven by condition (DPZ vs. PCB), timepoint (pre-vs. post-drug administration), and RSN (DAN, VAN, DMN, FPN). Specifically, we computed mean dual regression coefficients constrained within RSN-derived masks; subsequently, this data was included as the dependent variable in a LME^83^. Fixed effect predictors included condition, timepoint, RSN, and their interactions, with participant as a grouping factor. The model included participant-specific random intercepts and random slopes for condition and timepoint.

Planned contrasts interrogating drug-specific changes in RSN strength were computed as part of a LME on RSN strength ΔΔ. Drug-specific RSN change was computed as (post – pre)_DPZ_ – (post – pre)_PCB_ of mean regression coefficients. The LME formula was mean `RSN strength ΔΔ = RSN + (1 | participant)`. Estimated marginal means of strength were tested against a null hypothesis of ΔΔ = 0. To interrogate inter-network connectivity, Student’s one-sample t-testing was used with one-tailed alternative hypotheses.

To investigate drug effects on attentional orienting, we computed the DPZ–PCB difference in orienting index per-participant and examined the results using Student’s one-sample t-test. For the exploratory investigation of the relationship between attentional orienting improvement with DAN connectivity change, we first extracted mean dual regression coefficients from the cluster-wise-corrected DAN cluster mask, then computed the difference in coefficients between post-drug administration measurements (Δ-post). Each Δ-post input was correlated with change in orienting index using one-tailed Spearman’s correlation.

All statistical tests were performed in JASP (JASP Team 2025, Version 0.19.3). Default JASP model parameters were used unless otherwise stated. Where appropriate, multiple comparisons were Holm-corrected.

## Author contributions

PF and CS conceived the study. SH, LF, ES and PF collected the data. IG shared analysis tools. SH analysed the data and ran statistical analyses. SH, PF and CJS wrote the manuscript.

